# eDNA metabarcoding reveals a core and secondary diets of the greater horseshoe bat with strong spatio-temporal plasticity

**DOI:** 10.1101/2020.06.08.139584

**Authors:** Orianne Tournayre, Maxime Leuchtmann, Maxime Galan, Marine Trillat, Sylvain Piry, David Pinaud, Ondine Filippi-Codaccioni, Dominique Pontier, Nathalie Charbonnel

**Author notes:** equal contribution (Dominique Pontier, Nathalie Charbonnel).

## Abstract

Dietary plasticity is an important issue for conservation biology as it may be essential for species to cope with environmental changes. However, this process still remains scarcely addressed in the literature, potentially because diet studies have long been constrained by methodological limits. The advent of molecular approaches now makes it possible to get a precise picture of diet and its plasticity, even for endangered and elusive species. Here we focused on the greater horseshoe bat (*Rhinolophus ferrumequinum*) in Western France, where this insectivorous species has been classified as ‘Vulnerable’ on the Regional Red List in 2016. We applied an eDNA metabarcoding approach on 1986 fecal samples collected in six maternity colonies at three sampling dates. We described its diet and investigated whether the landscape surrounding colonies and the different phases of the maternity cycle influenced the diversity and the composition of this diet. We showed that *R. ferrumequinum* feed on a highly more diverse spectrum of prey than expected from previous studies, therefore highlighting how eDNA metabarcoding can help improving diet knowledge of a flying elusive endangered species. Our approach also revealed that *R. ferrumequinum* diet is composed of two distinct features: the core diet consisting in a few preferred taxa shared by all the colonies (25% of the occurrences) and the secondary diet consisting in numerous rare prey that were highly different between colonies and sampling dates (75% of the occurrences). Energetic needs and constraints associated with the greater horseshoe bat life-cycle, as well as insect phenology and landscape features, strongly influenced the diversity and composition of both the core and whole diets. Further research should now explore the relationships between *R. ferrumequinum* dietary plasticity and fitness, to better assess the impact of core prey decline on *R. ferrumequinum* populations viability.

## 1 INTRODUCTION

Food resources constitute a major environmental factor for animal populations (Hutchinson 1957; Schoener 1974). Both the quantity and quality of food resources are known to strongly impact the fitness of individuals (Eeva, Lehikoinen, et Pohjalainen 1997; Sorensen et al. 2009; Serrano-Davies et Sanz 2017) and in turn, the dynamics and viability of populations (Taylor et Schultz 2008; Vickery et al. 2001; Johnsen et al. 2017). Understanding species dietary requirements is therefore of main importance for designing conservation strategies (Cravens et al. 2018; Clare 2014; Brown et al. 2014; Groom et al. 2017). Several studies have even shown that some dietary characteristics could be related to elevated risk of species extinction. In particular, species at higher trophic levels in the food chain are more exposed to accumulative effects of pollutants along the food chain. This makes them more vulnerable to the deleterious effects of pollutants than species at lower trophic levels (Purvis et al. 2000; Careddu et al. 2015; Mann 2011). Besides, a narrow and specialized trophic niche (*i.e*. the range of possible prey) can also increase the vulnerability of species : specialists can face more constraints to respond to environmental changes in resource availability than generalist species (Boyles et Storm 2007; Pratchett, Wilson, et Baird 2006; Twining et al. 2019; Owens et Dittman 2003; Clavel, Julliard, et Devictor 2011). However, foraging can be a flexible activity. According to the optimal foraging theory, predators exploit resources that maximize the net energy intake whilst minimizing energetic costs through a trade-off between food profitability and searching time (Emlen 1966; MacArthur et Pianka 1966). As such, predators should be more selective on profitable resources when these latter are common, and more generalist when profitable resources are scarce. Such dietary plasticity is crucial to cope with environmental changes, including seasonal fluctuations (Bergmann et al. 2015; Kartzinel et Pringle 2015), climate change (de Oliveira et Val 2017; Durant et al. 2007) or anthropogenic pressure (Hempson et al. 2017; Quéméré et al. 2013; Smith et al. 2018). However, suboptimal diet can have negative impact on individual fitness (Sasakawa 2009). Dietary plasticity, despite its potential ecological and evolutionary importance, still remains scarcely addressed in the literature (Sousa, Silva, et Xavier 2019; but see Shutt et al. 2020), potentially because diet studies have long been constrained by methodological limits. Traditionally, diet has been described through direct observations of predatory events and/or through visual morphological analyses of prey remains in predator stomach, gut or scat (Nielsen et al. 2018; Sousa, Silva, et Xavier 2019). However, as these approaches depend on the presence of indigestible prey remains (e.g. exoskeletons), (i) prey are usually identified at a low taxonomic level (*i.e*. order or family ranks), (ii) soft-bodied prey are undetectable, and (iii) these approaches are very time consuming and are therefore usually limited in number of processed samples (Nielsen et al. 2018). The development of molecular approaches for identifying prey DNA contained in feces, in particular environmental DNA (eDNA) metabarcoding, has set aside most of the traditional methodologies limitations (Khanam et al. 2016; Clare 2014). Metabarcoding of fecal samples has been mainly applied for terrestrial predators, and more specifically for bats (Sousa, Silva, et Xavier 2019). Indeed these later are nocturnal, elusive, highly mobile and most of them are threatened, making it difficult to apply direct observations of their prey (Kunz et al. 2011; Kunz, Whitaker, and Wadanoli 1995; IUCN 2019). Insectivorous bats show a large range of foraging strategies, from specialists (e.g. mountain long-eared bat, *Plecotus macrobullaris;* Alberdi et al. 2012) to generalists (e.g. big brown bat *Eptesicus fuscus*; Clare, Symondson, et Fenton 2014). Bat species qualified as generalists can show preference towards certain prey (e.g. *Myotis daubentonii*; Vesterinen et al. 2016) and be more selective when their favorite prey are available in the environment (e.g. *Eptesicus fuscus*; Agosta, Morton, et Kuhn 2003).

The greater horseshoe bat (*Rhinolophus ferrumequinum*) is an insectivorous bat species whose diet and foraging behavior have been previously explored, especially in Northern Europe where it experienced severe declines this last century (Kervyn et al. 2009; Mathews et al. 2018; Pir 2009). These declines are likely due to anthropogenic factors (*i.e*. agricultural intensification or landscape urbanization) that might have resulted in losses of roosting habitats, habitat fragmentation and reduction in prey availability (Froidevaux et al. 2017; Mathews et al. 2018). *R. ferrumequinum* populations might therefore be affected by the global decrease of arthropods (Hallmann et al. 2017; Wickramasinghe et al. 2004; Strong et al. 1996). They might also be impacted by the accumulation of pesticides in body tissues which can decrease bat fitness (Bayat et al. 2014; Clarke-Wood et al. 2016; Stahlschmidt et Brühl 2012). *R. ferrumequinum* is even more vulnerable that it is a long-lived species (up to 30 years) with a low reproductive rate (maximum of one pup per year) and a late sexual maturity (two to five years) (Caubère, Gaucher, et Julien 1984; Ransome 1995; Wilkinson et South 2002).

Previous studies based on microscopy analyses have shown that *R. ferrumequinum* feeds mainly on three orders of arthropods : Lepidoptera, Coleoptera and Diptera (Jones 1990; Flanders and Jones 2009). They have also revealed that the proportion of each order of prey in the feces varied throughout the year with a preference for Lepidoptera when these latter were abundant in summer. An experimental study in controlled conditions showed that *R. ferrumequinum* can discriminate and select prey on the basis of the length and density of the prey in the environment, thanks to its very precise echolocation system (high and long constant-frequency enabling to compensate the Doppler shift) (Koselj, Schnitzler, et Siemers 2011). Altogether these studies suggested that *R. ferrumequinum* might have a plastic foraging strategy to maximize energy intake while minimizing energy costs (optimal foraging theory; Emlen 1966; MacArthur et Pianka 1966). Yet, energy cost could fluctuate between and within seasons as it has previously been shown for several bat species (e.g. summer peak of Myotis lucifugus energy expenditure during lactation; Kurta et al. 1989). In addition, *R. ferrumequinum* is very sensitive to the landscape surrounding the colonies, in particular to the vertical vegetation elements (Froidevaux et al. 2017; Pinaud et al. 2018; Wang et al. 2010), because of its short-distance echolocation system (up to 10m; Ortega, Moreno-Santillán, et Zamora-Gutierrez 2016). These features provide a large concentration and diversity of insects. They connect *R. ferrumequinum* foraging areas to colonies and they also provide protection against wind and predators (Verboom et Spoelstra 1999; Lewis 1969; Forman et Baudry 1984; Holland et Fahrig 2000). Acoustic and radio-tracking studies conducted around colonies (radius up to 10 kilometers) indicated that deciduous woodlands and pastures are preferential foraging areas for *R. ferrumequinum* (Flanders et Jones 2009; Jones 1990; Dietz, Pir, et Hillen 2013; Pinaud et al. 2018). Such landscapes are known to harbor rich insect communities, notably thanks to livestock and reduced soil modification. Yet, landscape effect on *R. ferrumequinum* diet and diet plasticity has never been explored. There is thus a growing need to simultaneously examine the temporal and spatial variations of *R. ferrumequinum* diet, but with much better taxonomic resolution and many more samples to overcome the detection and identification biases associated to the microscopy analyses. Such studies should enable us to better understand the influence of energetic constraints associated to the life cycle and the foraging landscape on dietary plasticity, thereby helping to improve the design of conservation strategies (e.g. preservation of key landscape and prey; Arrizabalaga-Escudero et al. 2015).

In this study, we analyzed the diet of *R. ferrumequinum* in Western France, an area with a strong responsibility for the conservation of this species (Vincent et Bat Group SFEPM 2014; Leuchtmann et al. 2019) classified as ‘Vulnerable’ on the Regional Red List. This area is dominated by an agricultural landscape and has experienced important changes in landscape features due to agricultural intensification since the 1960s (e.g. decrease of meadows, grasslands and hedges; increase in the average size of cultivated fields, increase in pesticides use) (Agreste 2016). We focused on the maternity season, as it corresponds to a period of high energy expenditure and foraging constraints for female bats due to gestation and lactation (Kurta et al. 1989; Mclean et Speakman 1999; Hughes et Rayner 1993; Henry et al. 2002). We specifically explored whether *R. ferrumequinum* diet varied between the different phases of the maternity cycle and/or were associated with landscape features surrounding the colonies. First, we expected that *R. ferrumequinum* diet should be more diverse in June and July - because of higher energy expenditures and constraints associated with the gestation and lactation - than in August when the young started to feed by themselves. We then hypothesized that a favorable environment (semi-open habitat composed of hedgerows and permanent meadows; Froidevaux et al. 2017; Flanders et Jones 2009) should favor a more diverse diet than a less favorable environment where foraging could be more constrained by local insect richness and profitability.

## 2 MATERIALS AND METHODS

### 2.1. Collection of guano samples

We collected 95 fecal pellets once a month from June to August 2018 beneath seven maternity colonies of *R. ferrumequinum* in Western France (Figure 1). The three sampling dates coincided with gestation (end of May to mid-June), lactation (mid-July to end of July) and post-lactation (mid-August to end of August) of *R. ferrumequinum*.

**Figure 1.**
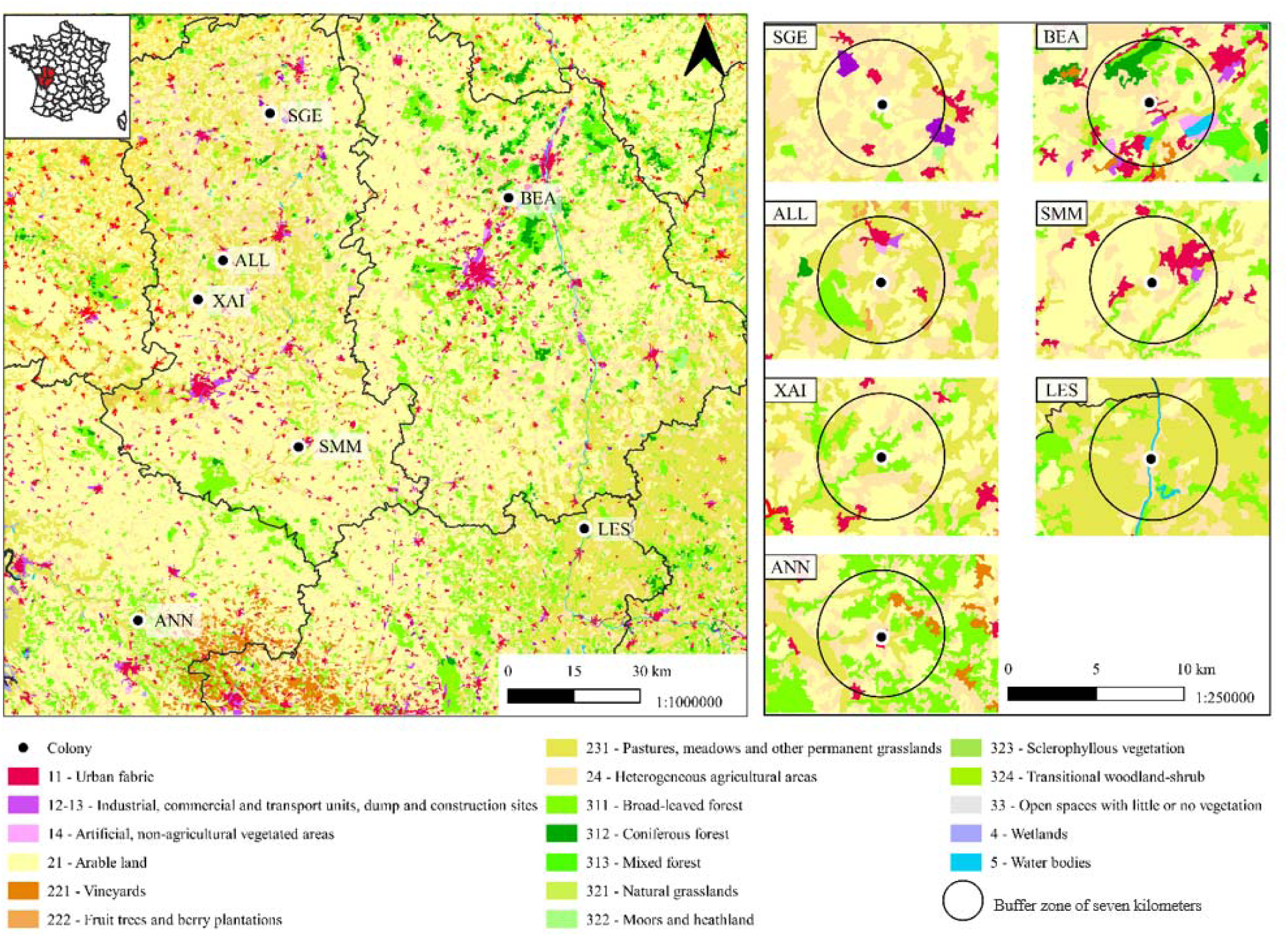
CORINE land cover map of the colonies included in this study: Allonne (ALL), Annepont (ANN), Beaumont (BEA), Lessac (LES), Sainte-Gemme (SGE), Saint-Martin-les-Melle (SMM) and Xaintray (XAI). Buffer zone of seven kilometers corresponds to the maximum hunting distance of *R. ferrumequinum* females during the lactation period.

As *R. ferrumequinum* often shares its maternity colonies with the Geoffroy’s bat (Myotis emarginatus) and as their droppings are undistinguishable, the high number of fecal samples collected ensured a sufficient number of *R. ferrumequinum* samples for diet analyses. Each pellet was retrieved from paper plates let on the ground during 10 days using single-usage pliers, and pellets were placed individually within a 96-well microplate. Paper plates were renewed at each collection date. This protocol enabled to minimize contaminations between samples. Samples were stored at −20°C until DNA extraction.

### 2.2. Characterization of the landscape surrounding bat colonies

We described landscape variables around *R. ferrumequinum* colonies within a buffer zone of seven kilometers, what corresponds to the average maximum hunting distance of *R. ferrumequinum* females during the lactation period (Pinaud et al. 2018). Data were extracted from the two French “Institut Géographique National” databases: “BDTopo” database (2018 version) for permanent vegetation (hedgerows, forests etc) and “Registre Parcellaire Graphique” (Graphic Parcel Register, GPR, 2017 version) for agricultural land use (crops). Hedge proportions corresponded to the sum of the hedgerow areas around the colonies. For GPR crops, the categories ‘Permanent meadow - predominant grass’ and ‘Long-rotation meadow (six years or more)’ were combined to form the variable ‘Permanent meadows’. The categories ‘Other temporary meadow of five years or less’ and ‘Ray-grass of five years or less’ were clustered into the variable ‘Temporary meadows’. Spatial operations on geographic entities were carried out using QGIS 3.4.6 Geographic Information System (QGIS Development Team 2019). We then applied a Principal Component Analysis (PCA) on these variables to characterize the landscape surrounding colonies (sum of the surface occupied in hectares). We used the *FactoMineR* R package to perform this analysis (Lê, Josse, et Husson 2008).

### 2.3. DNA extraction, PCR and library construction

DNA extraction was performed on pellet samples according to Zarzoso-Lacoste et al. (2018). Briefly, pellets were frozen at −80°C then bead-beaten for 2 x 30s at 30 Hz on a TissueLyser (Qiagen) using a 5-mm stainless steel bead. DNA was extracted using the NucleoSpin 8 Plant II kit (Macherey Nagel) with the slight modifications recommended in Zarzoso-Lacoste et al. (2018).

We amplified a 178-bp COI minibarcode using the primer set proposed by Vamos, Elbrecht, et Leese (2017) (fwhF1: YTCHACWAAYCAYAARGAYATYGG; fwhR1: ARTCARTTWCCRAAHCCHCC). These primers offer the best compromise to (1) maximize the detection and identification of arthropod prey and (2) identified the bat species to separate *R. ferrumequinum* and *M. emarginatus* diets and remove the pellets mixed by the DNA from these two bat species (Tournayre et al. in press). We used the metabarcoding protocol described in Galan et al. (2018) with the PCR programs optimized in Tournayre et al. (in press). We included in each 96-well microplate a negative control for extraction (NC_ext_), a negative control for PCR (NC_PCR_) and a negative control for indexing (NC_index_). All the DNA extractions and controls were analyzed by three independent technical replicates, *i.e*. three sequencing libraries using different dual-index. The PCR_1_, corresponding to the gene specific amplification, was performed in 10 µl reaction volume using 5 µl of 2x Qiagen Multiplex Kit Master Mix (Qiagen), 2.5 µl of ultrapure water, 0.5 µl of each mix of forward and reverse primers (10 µM), and 1.5 µl of DNA extract. The PCR_1_ conditions consisted in an initial denaturation step at 95°C for 15 min, followed by 40 cycles of denaturation at 94°C for 30 s, annealing at 45°C for 45 s, and extension at 72°C for 2min, followed by a final extension step at 72°C for 10 min. The PCR_2_ consisted in a limited-cycle amplification step to add multiplexing index i5 and i7 (8 bases each) and Illumina sequencing adapters P5 and P7 at both ends of each DNA fragment from PCR_1_. PCR_2_ was carried out in a 10µl reaction volume using 5 µl of Qiagen Multiplex Kit Master Mix (Qiagen) and 2µL of each indexed primer i5 and i7 (0.7 µM). Then, 2 µL of PCR_1_ product was added to each well. The PCR_2_, corresponding to the sample-specific dual indexing, started by an initial denaturation step of 95°C for 15 min, followed by 8 cycles of denaturation at 94°C for 40 s, annealing at 55°C for 45 s and extension at 72°C for 2min followed by a final extension step at 72°C for 10 min. PCR_2_ products were pooled and put on a low-melting agarose gel (1.25%) for excision. We used the PCR Clean-up Gel Extraction kit (Macherey-Nagel) to purify the excised bands. DNA pool was quantified using the KAPA library quantification kit (KAPA Biosystems), normalized at 4nM before loading 14 pM and 5% of PhiX control on a MiSeq flow cell with a 500-cycle Reagent Kit v2 (Illumina). The samples were sequenced on five Illumina runs.

### 2.4. Bioinformatics and taxonomic assignments

We used the R pre-processing script from Sow et al. (2019) to merge pair sequences into contigs using FLASH v.1.2.11 (Magoč et Salzberg 2011) and to trim primers using CUTADAPT v.1.9.1 (Martin 2011). We then used FROGS pipeline (‘Find Rapidly OTU with Galaxy Solution’, Escudié et al., 2018) to create an abundance table for each Operational Taxonomic Units (OTUs). FROGS pipeline enabled to (i) filter sequences by length (+/- 20 bp from the expected length), (ii) cluster in very resolutive OTUs the sequences using a maximum aggregation distance of one mutation with the SWARM algorithm (Mahé et al. 2014), (iii) remove chimeric sequences using VSEARCH with *de novo* UCHIME method (Edgar et al. 2011; Rognes et al. 2016), and (iv) filter by keeping only OTUs present in at least two libraries. In addition we used *isBimeraDenovo* from *dada2* (Callahan et al. 2016) to remove the residual chimeric sequences which were not detected using FROGS pipeline. We also used the T_CC_ and T_FA_ thresholds approach proposed by Galan et al. (2016) to filter cross-contaminations during the laboratory procedure and to filter the false assignments of reads to a PCR product due to the generation of mixed clusters during the sequencing, respectively. Lastly, we considered that a sample was positive for a particular OTU if all replicates (3 out of 3) were positive for this taxon. This procedure enabled to remove inconsistent OTUs due to PCR or sequencing errors, what reduced the number of putative false-positive results (Alberdi et al., 2018; Tournayre et al. in press). Finally, for each sample and OTU, the reads obtained for the three PCR replicates were summed.

Taxonomic assignments were carried out using the NCBI BLAST+ automatic affiliation tool available in FROGS pipeline, with the Public Record Barcode Database (data related to BOLD database http://v3.boldsystems.org in February 2019, with maximum 1% of N). Arthropod species that were not referenced in Europe according to Fauna Europea (Jong et al. 2014)or INPN (Muséum national d’Histoire naturelle, 2003), as well the microscopic species (e.g. Macrochelidae, Uropodidae, Cheyletidae), were discarded from the dataset. Finally, among the identified prey, we looked for agricultural pests according to the Arthemis database (http://arthemisdb.supagro.inra.fr/; 2,185 species of insects listed as pests on the date of database extraction - 04/10/2019).

### 2.5. Diet analyses

Diet analyses were only conducted on *R. ferrumequinum* samples that were not contaminated by other vertebrate species (other bats, birds or rodents).

#### 2.5.1. Reliability of the data

We checked for appropriate sequencing depth per sample to ensure reliable comparisons across samples using the function *depth.cov* from the R package *hilldiv* (Alberdi et Gilbert 2019). This function gives the percentage of estimated diversity covered in each sample. Then, we assessed our efficiency in describing prey diversity with respect to sampling effort by generating taxa accumulation curves for each colony, sampling date and considering each taxonomic rank (order, family, genus, species). We used the function *specaccum* of the R package *vegan* (Oksanen et al. 2019) and we applied 1,000 random re-sampling events. As many prey may remain undetected, we used the function *specpool* of the package *vegan* to estimate the Chao index (Chao 1987), which gave the total estimated richness (*i.e*. both undetected and observed prey).

#### 2.5.2. Diet diversity and composition analyses

##### Alpha diversity

Alpha diversity analyses were carried out using Hill numbers from the R package *hilldiv*. Hill numbers enable to modulate the relative weight of abundant and rare OTUs through a single parameter *q* (the order of diversity). Alpha diversities were computed for (i) *q*⍰=⍰0 (it corresponds to prey richness; the same weight is attributed to all OTUs) and (ii) *q*⍰=⍰1 (it corresponds to Shannon diversity that considers both richness and evenness). For each level of taxonomic resolution, we tested the effects of sampling date, landscape (described as the coordinates of the colonies on the main axes of the PCA) and their interaction on alpha diversity using quasi-Poisson Generalized Linear Models (GLMs). We used the false discovery rate (FDR) to account for multiple testing (Benjamini et Hochberg 1995). The adjusted p-value thresholds after FDR correction (*p*_critical_) were estimated following Castro & Singer (2006).

##### Beta diversity

The spatio-temporal variation of *R. ferrumequinum* diet composition was first explored using histograms of the frequency of prey occurrence, computed with the ggplot2 R package (Wickham 2016). For each level of taxonomic resolution, we applied the permutational multivariate analysis of variance (perMANOVA; 999 permutations) using the *adonis* (*Analysis of variance using distance matrices*) function from the R package *vegan* (Oksanen et al. 2019) to investigate whether diet composition differed between sampling dates and colonies. Homogeneity of variance was tested using the betadisper function of the package *vegan*. The adjusted *p*-value thresholds after FDR correction (*p*_critical_) were estimated following Castro & Singer (2006).

Dissimilarities between prey communities (Bray-Curtis matrix) were visualized using the Nonmetric Multidimensional Scaling (NMDS) with the *metaMDS* function of the R package *vegan* (try = 20, trymax = 5,000). The quality of the solution was evaluated based on the stress value: stress values lower than 0.05 indicate that the solution is of excellent quality and stress values higher than 0.2 indicate solution of poor quality (Kruskal 1964).

Finally, as the landscape could influence the prevalence of taxa in the environment (e.g. presence or absence of prey ecologic requirements), we investigated whether diet composition between colonies were correlated with landscape composition using Mantel test (999 permutations) between diet and landscape dissimilarity matrices. Mantel tests were computed using the mantel.rtest function from the R package ade4 (Dray et Dufour 2007).

## 3 RESULTS

### 3.1 Pre-treatment of high-throughput sequencing data

#### 3.1.1. Data filtering

We obtained 26,737 OTUs (44,801,072 sequences) after applying FROGS pipeline. 8,839 supplementary OTUs were ruled out because they were considered as chimera by *iBimeraDenovo* tools from *dada2* (33.05% of the OTUs). After filtering using the controls and triplicates, we obtained 7,103 OTUs (41,216,468 sequences) for 1,986 samples. Over the 7,103 OTUs 2,110 could not be taxonomically assigned due to low confidence level (< 97% identity and/or < 90% coverage) and 3,204 were absent from database. Among the 1,178 genuine assigned OTUs, 72 OTUs were discarded because they were not referenced in Europe; they were assigned by error to insect species in BOLD but correspond to bacteria (e.g. *Wolbachia*, Rickettsiales) or they were likely to be environmental contaminations (e.g. mites).

#### 3.1.2 Identification of the predator

Among the 1,986 samples obtained after the filtering steps, 1,194 corresponded to *R. ferrumequinum* (60.1%), 381 to *M. emarginatus* (19.2%), 32 to the greater mouse-eared bat (Myotis myotis; 1.6%), one to the genus Serotinus (0.05%). 29 samples were identified as a mix of two bat species, including *R. ferrumequinum, M. emarginatus, Serotinus* sp., the long-eared bat (*Plecotus auritus*), the Bechstein’s bat (*Myotis bechsteinii*) and the Natterer’s bat (*Myotis nattereri*). Twelve samples were identified as rodents or a mix of rodents and bats (0.6%), 15 samples as birds (*Hirundo* sp.; 0.7%) and 91 samples as a mix of birds and bats (4.5%). Finally, we could not identify any predator for 178 samples (8.9%) and neither predator nor prey identification for 52 samples (2.6%), probably because OTUs of these samples were below the filtering thresholds used to clean the data (*i.e*. false negatives).

### 3.2 Landscape characterization

The colony of Saint-Martin-les-Melle has been ruled out from the analyses because of an insufficient number of *R. ferrumequinum* samples in June (*N* = 0) and in July (*N* = 1). Then, 82 samples among the 1,115 from the six remaining colonies were discarded because of an absence of prey detection after all the previous filters.

Principal components analysis of the sum of the vegetation surface surrounding the colonies (ha) clearly separated two types of landscapes with the first two PCA axes explaining the largest part of the total inertia. The first axis (Axis.1, 48.34%) represented a landscape gradient running from a habitat dominated by forests (Beaumont) to a semi-open habitat dominated by meadows and hedgerows (Lessac, Sainte-Gemme, Allonne, Xaintray), with an intermediate situation in Annepont (Figure 2). The second axis (Axis.2, 25.90%) separated the colony of Annepont - characterized by vineyards and deciduous forests - from all other colonies (Figure 2).

**Figure 2.**
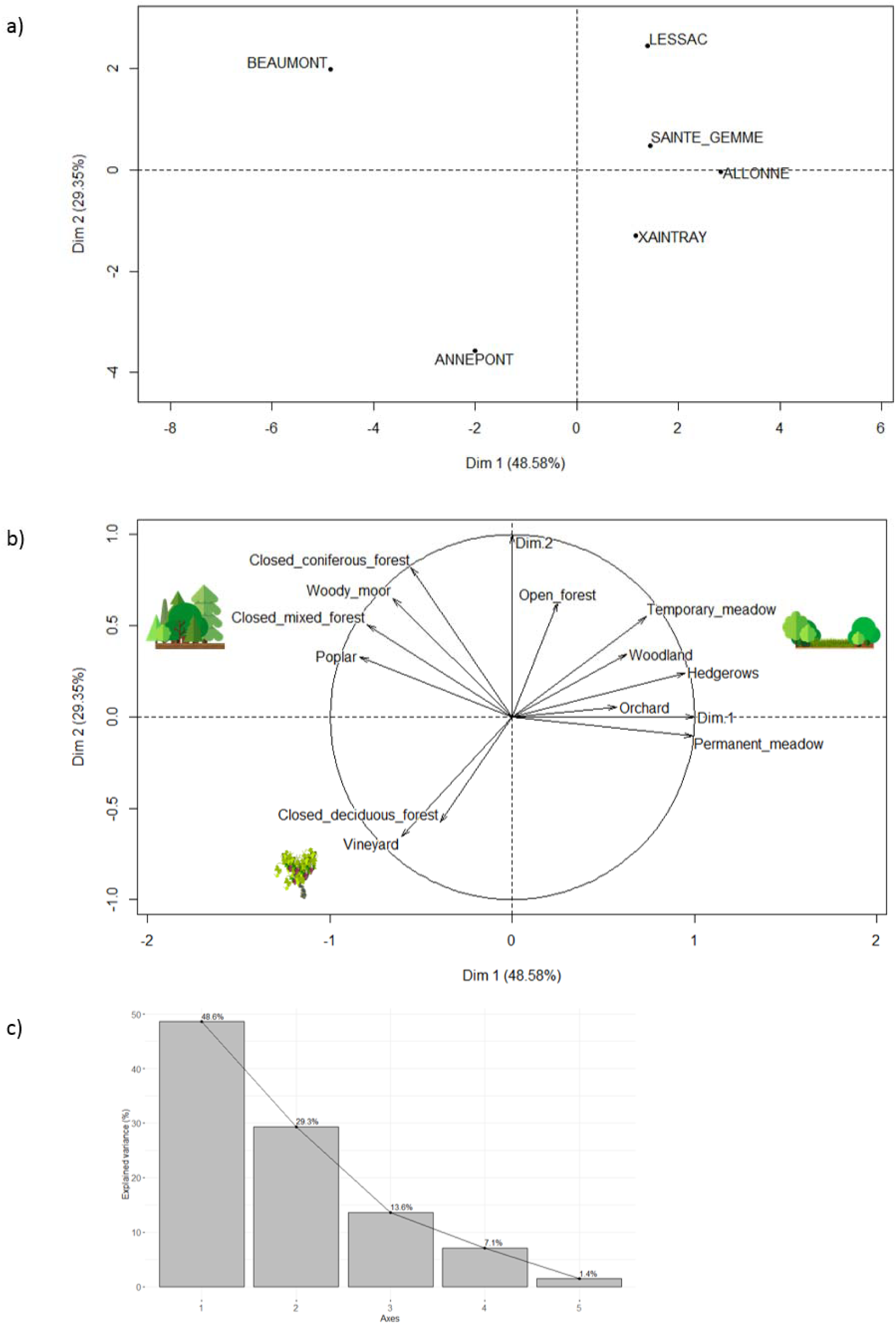
Principal Component Analysis (PCA) of twelve features describing the landscape surrounding the colonies (radius = 7km, surface in hectare). Representation of a) the six colonies, b) the landscape variables and c) the eigenvalue graph which indicates the percentage of variance explained by each axis of the PCA.

### 3.3 Diet analyses

#### 3.3.1. Reliability of the data

Our results revealed an appropriate sequencing depth per sample, with the estimated diversity covered in each sample at *q* = 0 (*i.e*. richness) and *q* = 1 (*i.e*. Shannon diversity; both richness and evenness) ranging between 95% and 100%.

The accumulation curves reached the plateau at the order level, with Chao index values being very close to the number of taxa observed in all colonies and sampling dates (Figure S1). However, the more precise the taxonomic level was, the more insufficient was the sampling effort to recover all the diversity. At the family, genus and species levels, the plateau was not reached and some strong differences between Chao index values and the numbers of observed taxa were observed (Figure S1).

#### 3.3.2. Variability of the whole and core diets

The final complete dataset included 1,033 *R. ferrumequinum* samples corresponding to six colonies and collected at three sampling dates. We identified 679 taxa from 17 arthropod orders, 124 families, 434 genera and 559 species (Figure 3). Three main orders were detected: Lepidoptera (57% of the occurrences), Diptera (23%) and Coleoptera (13%). We identified an important number of agricultural pests with 133 species notified in the Arthemis database (Table S1) and representing 31.86% of the occurrences. Pest species were mainly Lepidoptera (70.9%; e.g. *Thaumetopoea pityocampa, Archips podana*), Coleoptera (13.9%; e.g. *Curculio elephas, Melolontha melolontha*), Diptera (12.2%; e.g. *Tipula lateralis, Nephrotoma appendiculata*) and Hemiptera (2.7%; e.g. *Adelphocoris lineolatus, Fieberiella florii*).

**Figure 3.**
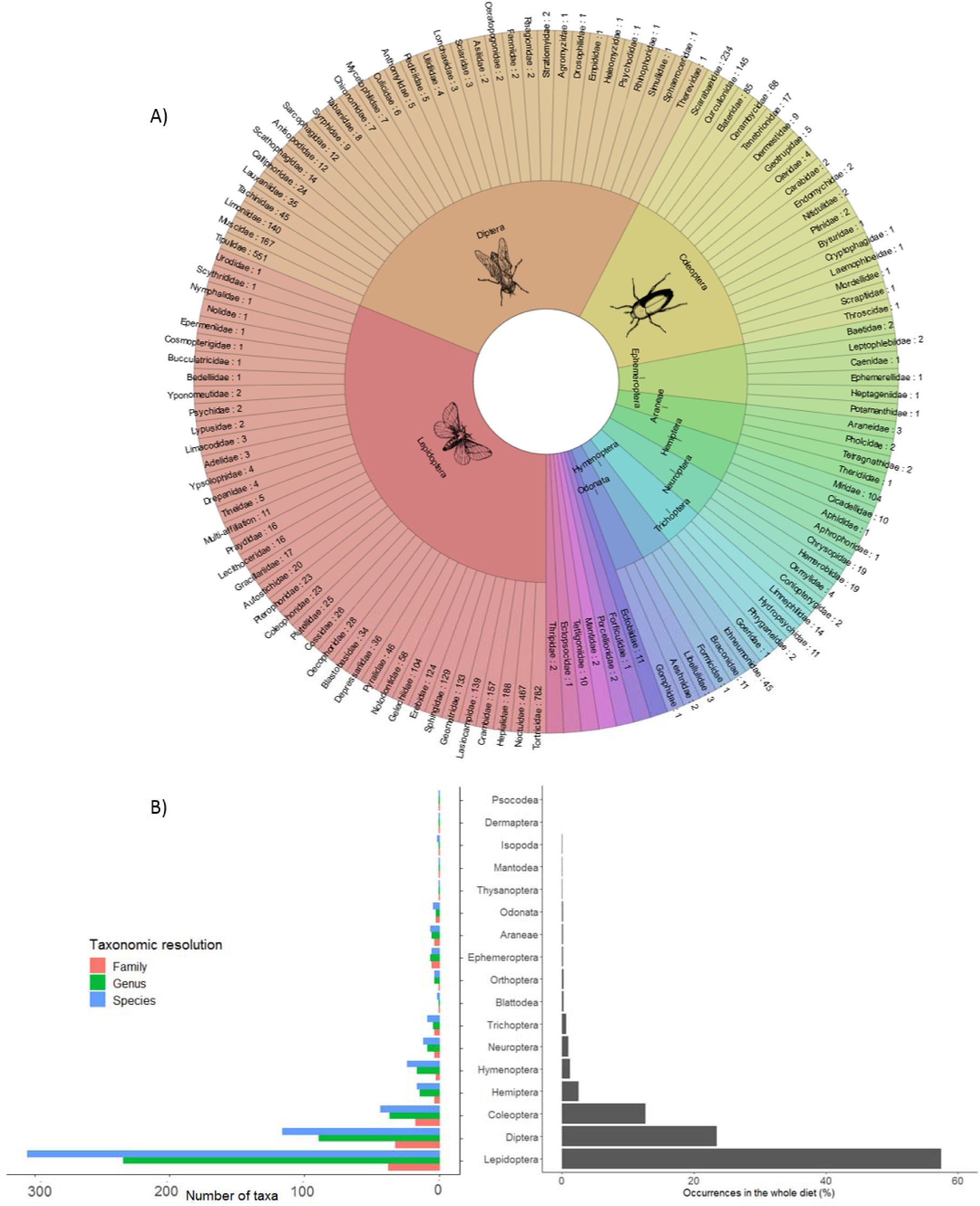
Representation of the prey taxonomic diversity detected from *R. ferrumequinum* guano, considering six colonies and three sampling dates. A) The two circles represent taxonomic ranks from the families outwards to the order in the center. Numbers correspond to the number of occurrences of each taxonomic family in the whole diet. B) Taxonomic diversity of the arthropod orders expressed as the number of prey families, genus, species (left) and percentage of occurrences (right) in the whole diet.

We also noticed that this complete dataset included a large proportion of rare prey. 59% of prey taxa were represented by only two (103 taxa) or even only one (300 taxa) occurrence over the 1,033 samples. We therefore considered a smaller dataset (hereafter called ‘core diet’) composed of the most frequent prey species (frequencies of occurrences > 5%). They represented 2.50% of all the taxa detected over the 1,033 samples, 24.89% of the occurrences and they accounted for 46.26% of the total number of reads. In this core diet, we identified 17 taxa from three orders (Lepidoptera, Diptera, Coleoptera), 10 families, 14 genera and 15 species (Figure 4). The 15 species - from the most occurrent (106 occurrences) to the less occurrent (52 occurrences)- were: *Celypha striana* (Lepidoptera), *Serica brunnea* (Coleoptera), *Limonia nubeculosa* (Diptera), *Musca autumnalis* (Diptera), *Triodia sylvina* (Lepidoptera), *Tortrix viridana* (Lepidoptera), *Laothoe populi* (Lepidoptera), *Stenagostus rhombeus* (Coleoptera), *Tipula fascipennis* (Diptera), *Korscheltellus lupulina* (Lepidoptera), *Agrotis bigramma* (Lepidoptera), *Copris lunaris* (Coleoptera), *Zeiraphera isertana* (Lepidoptera), *Tipula maxima* (Diptera) and *Euthrix potatoria* (Lepidoptera) (Table S2). Three agricultural pest species and one vector of livestock disease agents were identified in this core diet: *S. brunnea, T. viridana, T. maxima* and *M. autumnalis*, respectively.

**Figure 4.**
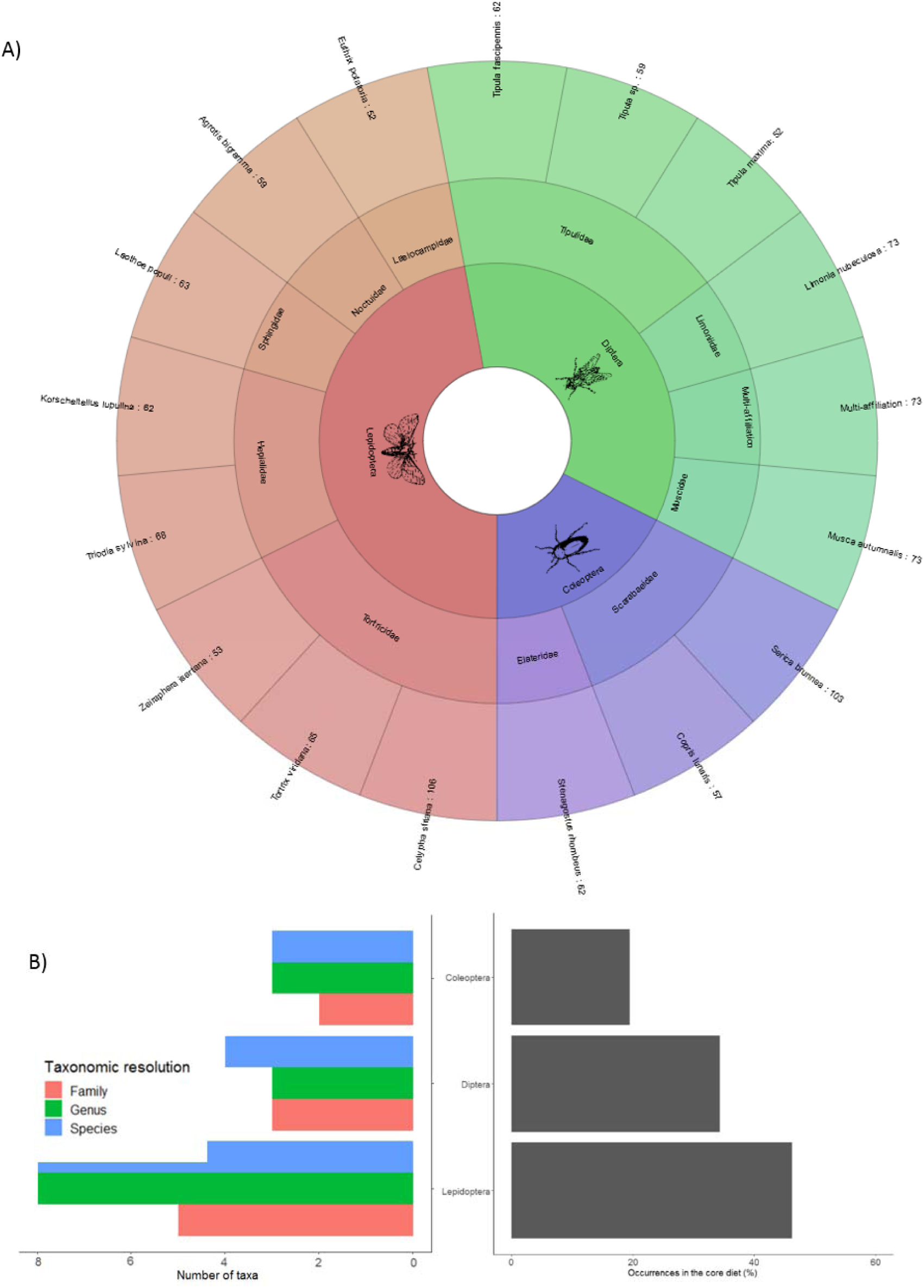
Representation of the prey taxonomic diversity detected from *R. ferrumequinum* guano, considering six colonies and three sampling dates and considering only the prey with an occurrence frequency > 5%. A) The circles represent taxonomic ranks from the species outwards to the order in the center. B) Relative representation of the frequent prey expressed as the number of prey families, genus, species (left) and percentage of occurrences (right).

Based on this result, we defined the core diet as all the prey taxa with more than 5% of occurrences frequency as the core diet. The whole diet corresponds to the core diet plus the secondary diet, *i.e*. the prey taxa with less than 5% of occurrence frequency. Further analyses of alpha and beta diversity were performed both on the complete (whole diet) and restricted (core diet) datasets.

#### 3.3.2. Alpha diversity

Considering the whole diet, we found a significant effect of the first PCA axis (Axis.1) and of the sampling date (month) whatever the taxonomic level and *q* value considered.

The interaction between Axis.1 and sampling date was significant for *q* = 0 and all taxonomic levels except family, and for *q* = 1 and the order level only (Table 1, Figure S2). Alpha diversity increased along Axis.1 in June and July, then slightly decreased in August at the order, family and genus level but remained stable at the species level (Table 1, Figure 5A). Alpha diversity remained slightly equivalent throughout the summer in the two colonies located in landscape dominated by forests (left of Axis.1, Figure 5A) while it was lower in August in the four colonies located in landscape dominated by meadows and hedgerows (right of the Axis.1, Figure 5A). At the family (*q* = 0 and *q* = 1), genus (*q* = 1) and species (*q* = 1) levels, we observed a positive effect of Axis.1 on alpha diversity and a peak of alpha diversity in July (Table 1, Figures 5B).

**Table 1.** GLMs results testing the effect of landscape variables (Axis 1 and 2 of the PCA) and sampling months (June, July, August) on the estimates of alpha diversity for *q* = 0 and *q* = 1 at each taxonomic level: order, family, genus and species. Significant *p*-values after correction for FDR multiple tests are represented in bold. A) On the whole diet, B) on the core diet.

| A) | | $q = 0$ | | | $q = 1$ | | |
| --- | --- | --- | --- | --- | --- | --- | --- |
| | | Df | F | $p$ | Df | F | $p$ |
| Order | Axis 1 | 1 | 18.040 | **2.360e-05** | 1 | 14.322 | **0.0002** |
|  | Month | 2 | 35.875 | **8.7290e-16** | 2 | 25.973 | **1.077e-11** |
|  | Axis 2 | 1 | 4.191 | 0.041 | 1 | 4.190 | 0.041 |
|  | Axis 1 * Month | 2 | 15.771 | **1.796e-07** | 2 | 12.853 | **3.132e-06** |
|  | Axis 2 * Month | 2 | 0.131 | 0.877 | 2 | 0.0004 | 0.999 |
| Family | Axis 1 | 1 | 18.001 | **2.408e-05** | 1 | 16.603 | **5.015e-05** |
|  | Month | 2 | 25.019 | **2.466e-11** | 2 | 30.548 | **1.457e-13** |
|  | Axis 2 | 1 | 0.548 | 0.459 | 1 | 1.944 | 0.164 |
|  | Axis 1 * Month | 2 | 4.383 | 0.013 | 2 | 1.814 | 0.164 |
|  | Axis 2 * Month | 2 | 1.133 | 0.323 | 2 | 0.962 | 0.383 |
| Genus | Axis 1 | 1 | 19.247 | **1.269e-05** | 1 | 15.919 | **7.151e-05** |
|  | Month | 2 | 12.139 | **6.166e-06** | 2 | 25.249 | **2.152e-11** |
|  | Axis 2 | 1 | 0.065 | 0.799 | 1 | 1.693 | 0.193 |
|  | Axis 1 * Month | 2 | 8.588 | **0.0002** | 2 | 3.055 | 0.048 |
|  | Axis 2 * Month | 2 | 4.232 | 0.015 | 2 | 2.156 | 0.116 |
| Species | Axis 1 | 1 | 23.805 | **1.238e-06** | 1 | 16.593 | **5.048e-05** |
|  | Month | 2 | 10.608 | **2.759e-05** | 2 | 17.081 | **5.259e-08** |
|  | Axis 2 | 1 | 0.169 | 0.681 | 1 | 1.503 | 0.220 |
|  | Axis 1 * Month | 2 | 6.452 | **0.002** | 2 | 2.179 | 0.114 |
|  | Axis 2 * Month | 2 | 3.864 | 0.021 | 2 | 1.670 | 0.189 |
| B) | | $q = 0$ | | | $q = 1$ | | |
| | | Df | F | $p$ | Df | F | $p$ |
| Order | Axis 1 | 1 | 19.486 | **1.173e-05** | 1 | 17.848 | **2.743e-05** |
|  | Month | 2 | 21.638 | **7.617e-10** | 2 | 13.346 | **2.102e-06** |
|  | Axis 2 | 1 | 0.107 | 0.744 | 1 | 0.110 | 0.741 |
|  | Axis 1 * Month | 2 | 2.266 | 0.104 | 2 | 1.265 | 0.283 |
|  | Axis 2 * Month | 2 | 3.592 | 0.028 | 2 | 0.846 | 0.429 |
| Family | Axis 1 | 1 | 38.443 | **9.663e-10** | 1 | 40.056 | **4.686e-10** |
|  | Month | 2 | 13.463 | **1.834e-06** | 2 | 10.945 | **2.126e-05** |
|  | Axis 2 | 1 | 1.278 | 0.259 | 1 | 1.918 | 0.166 |
|  | Axis 1 * Month | 2 | 4.112 | 0.017 | 2 | 1.513 | 0.221 |
|  | Axis 2 * Month | 2 | 0.642 | 0.526 | 2 | 0.178 | 0.837 |
| Genus | Axis 1 | 1 | 36.904 | **2.042e-09** | 1 | 37.617 | **1.521e-09** |
|  | Month | 2 | 15.794 | **1.958e-07** | 2 | 11.086 | **1.854e-05** |
|  | Axis 2 | 1 | 2.227 | 0.136 | 1 | 3.221 | 0.073 |
|  | Axis 1 * Month | 2 | 4.022 | 0.018 | 2 | 1.451 | 0.235 |
|  | Axis 2 * Month | 2 | 0.989 | 0.372 | 2 | 0.002 | 0.998 |
| Species | Axis 1 | 1 | 32.795 | **1.533e-08** | 1 | 33.239 | **1.286e-08** |
|  | Month | 2 | 12.358 | **5.338e-06** | 2 | 7.265 | **0.0007** |
|  | Axis 2 | 1 | 0.461 | 0.497 | 1 | 0.400 | 0.527 |
|  | Axis 1 * Month | 2 | 3.781 | 0.023 | 2 | 1.246 | 0.288 |
|  | Axis 2 * Month | 2 | 1.943 | 0.144 | 2 | 0.468 | 0.626 |

**Figure 5.**
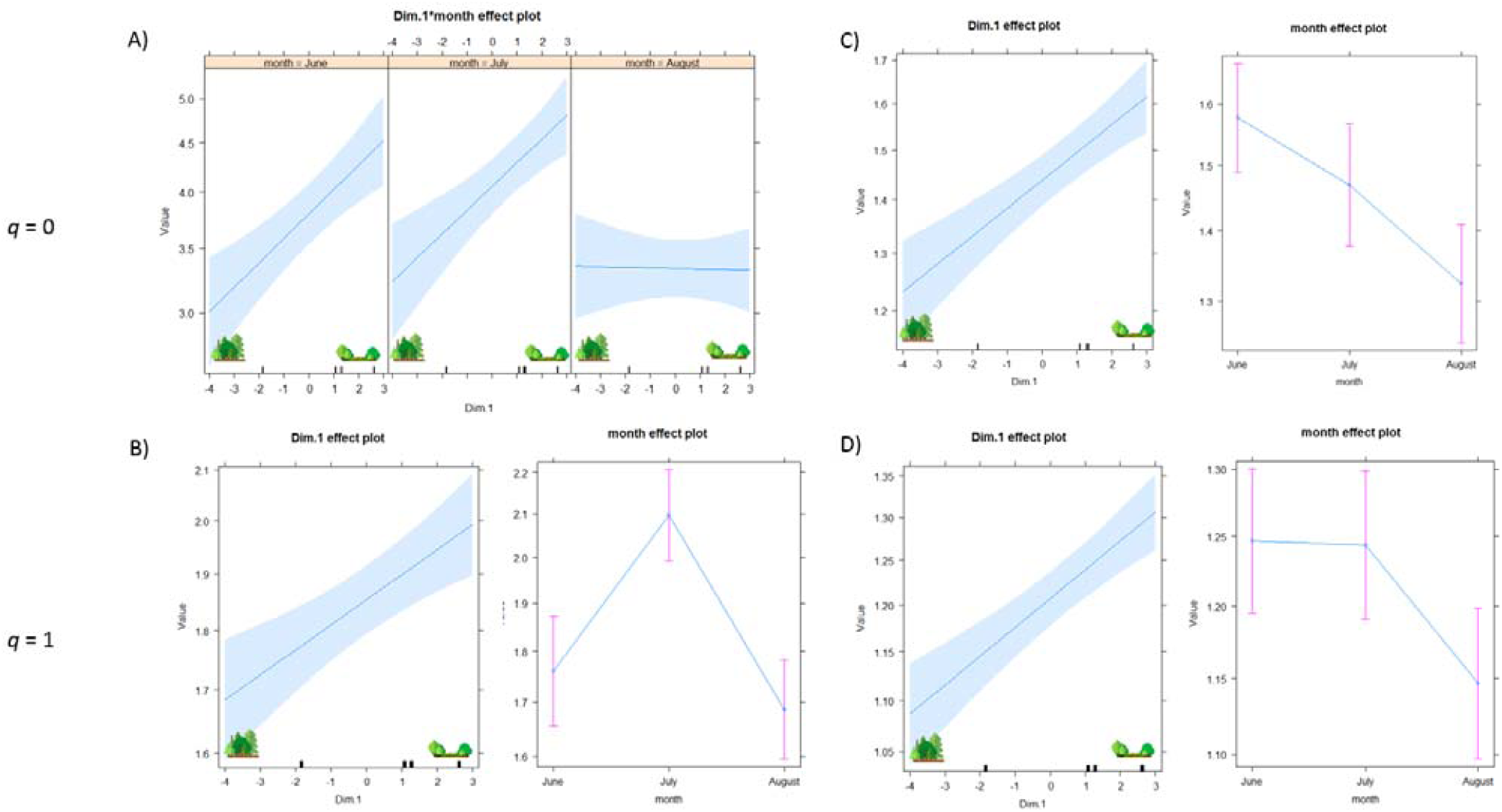
Plot of the variable effects on alpha diversity estimates at the species level: A) the whole diet and *q* = 0, B) the whole diet and *q* = 1, C) the core diet and *q* = 0 and D) the core diet and *q* = 1. ‘Dim.1’ represented Axis.1 of the PCA and ‘month’ the sampling date effect (June, July and August). The dominant landscapes - as determined by the axis 1 of the PCA – are indicated.

Considering the core diet, the effects of sampling date and landscape (Axis.1) were always significant whatever the taxonomic levels and *q* values considered (Table 1). We observed a negative effect of sampling date and a positive effect of Axis.1 on alpha diversity (Figures 5C and 5D).

All results are detailed in Table 1 and Figure S2.

#### 3.3.3. Beta diversity

Considering the whole diet, the relative occurrence of Lepidoptera and Diptera showed opposite patterns of temporal variations with an increase and a decrease over the summer, respectively (Figure 6A). The core diet showed different patterns. The colonies of Annepont (landscape dominated by vineyards and forest), Lessac, Allonne, Sainte-Gemme and Xaintray (all characterized by hedgerows and meadows) exhibited a peak of Diptera and a minimum of Lepidoptera in July (Figure 6B). The colony of Beaumont (landscape dominated by forests) showed a singular pattern: whatever the sampling date considered, the prey community was dominated by a single order (Coleoptera in June, Lepidoptera in July and August) and a maximum of two species (Figure 6B).

**Figure 6.**
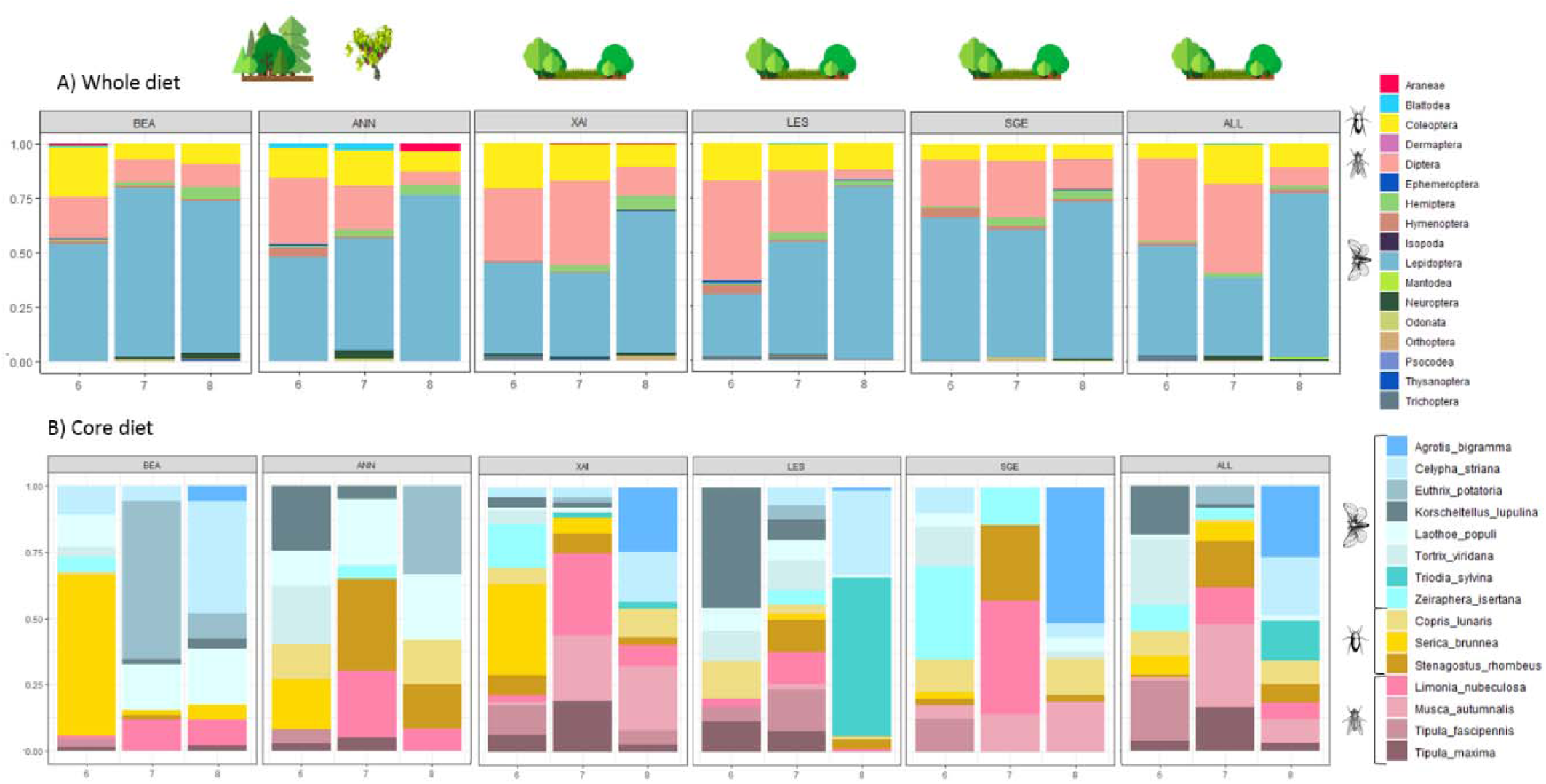
Percentage of prey occurrence in each of the six colonies - sorted according to their coordinates on the Axis.1 of the PCA (Allonne ‘ALL’, Annepont ‘ANN’, Beaumont ‘BEA’, Lessac ‘LES’, Sainte-Gemme ‘SGE’, Xaintray ‘XAI’) - for the three dates of sampling (‘6’ = June, ‘7’ = July, ‘8’ = August). A) The whole diet considered at the order level of taxonomic resolution. B) The core diet considered at the species level of taxonomic resolution. In both figures, the three most occurrent orders (Lepidoptera, Diptera, Coleoptera) are represented by an icon. The dominant landscapes - as determined by the axes 1 and 2 of the PCA - are indicated for each colony.

PerMANOVA analyses revealed significant variations of *R. ferrumequinum* diet between dates of sampling and colonies, for both the whole and core diets (Table 2). Note however that the variance between groups (colonies or sampling dates) were not homogeneous (Table 2). NMDS plots built on the whole diet highlighted the presence of rare prey that exacerbated the dissimilarity between colonies and sampling dates (Figures 7A and 7B). When considering the core diet, NMDS plots showed a high overlap of prey between sampling dates and colonies (Figures 7C and 7D). At the species level, NMDS plots revealed a high overlap between colonies but also suggested a pattern corresponding to the summer progression (from June to August), with less prey species shared between June (e.g. *Tortix viridana, Zeiraphera isertana*) and August (e.g. *Triodia sylvina, Agrotis bigramma*) (Figure 7D).

**Table 2.**
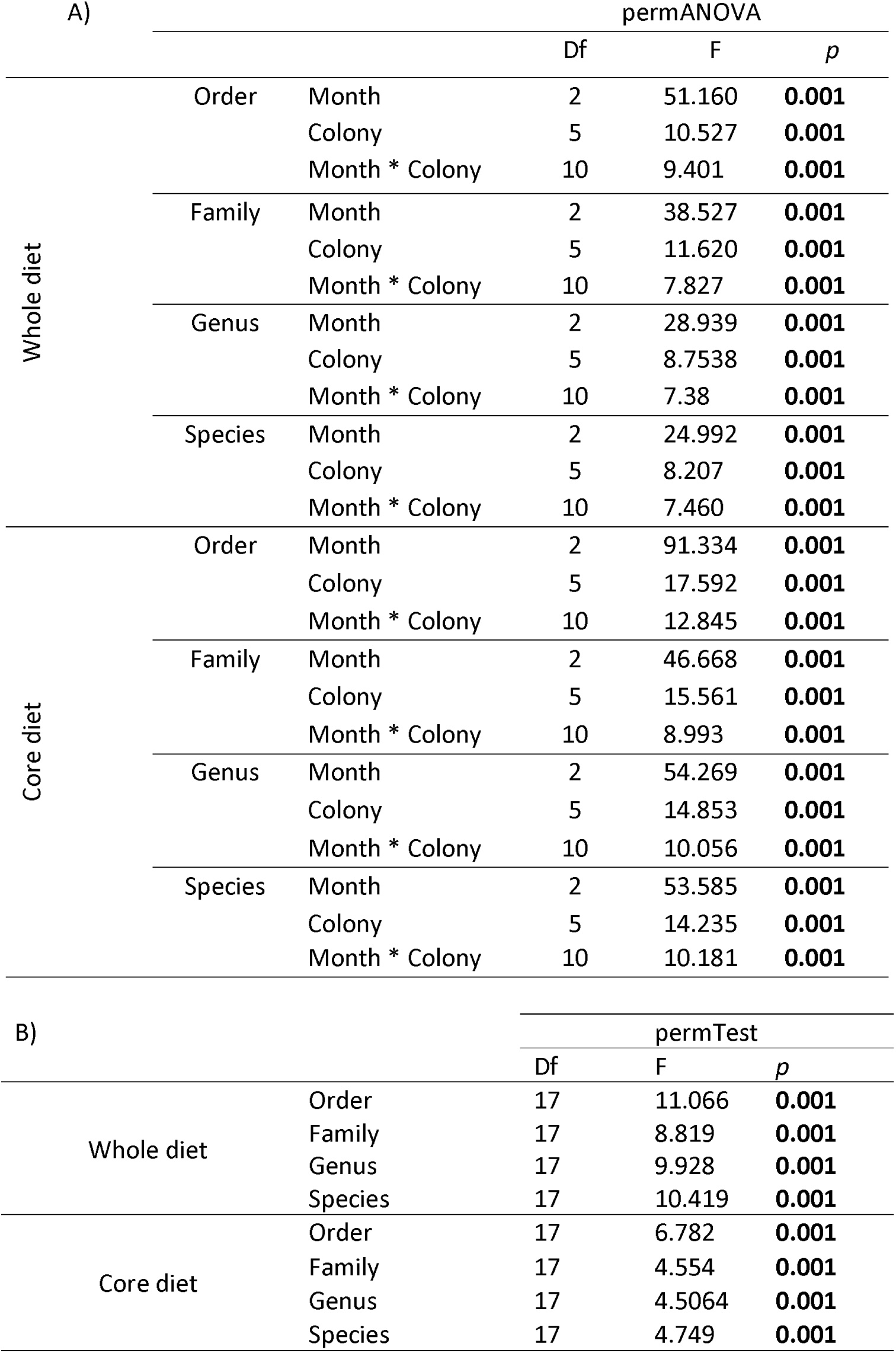
Results of A) permANOVA analyses and B) homogeneity of variance analysis by permutation (999 permutations), based on the Bray-Curtis dissimilarity for each taxonomic rank tested. Significant *p*-values after correction for multiple FDR tests are represented in bold. These analyses were performed on the whole diet and on the core diet.

**Figure 7.**
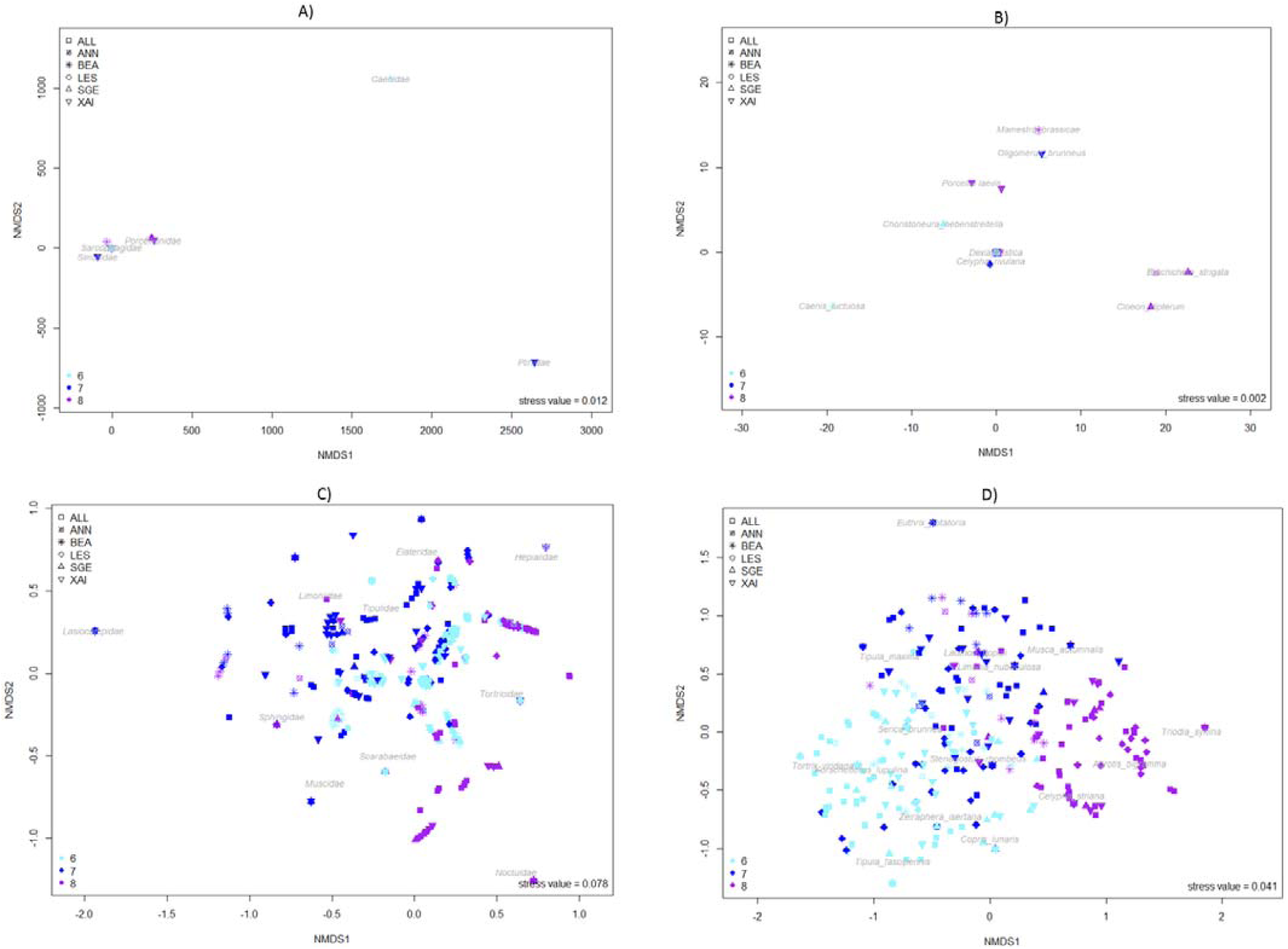
Representation of the Bray-Curtis dissimilarity using NMDS. A) Whole diet at the family level of taxonomic resolution, B) Whole diet at the species level of taxonomic resolution, C) Core diet at the family level of taxonomic resolution, D) Core diet at the species level of taxonomic resolution. Abbreviation: 6 = June (cyan), 7 = July (blue), 8 = August (purple), ANN = Annepont (circle cross), ALL = Allonne (square), BEA = Beaumont (star), LES = Lessac (diamond), SGE = Sainte-Gemme (triangle point up) and XAI = Xaintray (triangle point down).

Finally, we observed a positive relationship between the beta diversity (differentiation between diet composition) and the dissimilarity of the landscape surrounding the colonies when considering the whole diet (0.002 < *p* < 0.006) or the core diet (0.001 < *p* < 0.029) at all levels of taxonomic resolution, except at the order level for the whole diet (*p* = 0.478).

## 4 DISCUSSION

Based on eDNA metabarcoding, our study showed that *R. ferrumequinum* diet is much more diversified than previously described by microscopic (Jones 1990; Flanders et Jones 2009) and molecular analyses (Galan et al. 2018; Alberdi et al. 2020; Aldasoro et al. 2019). We revealed a broader ecological and taxonomic variety of prey from the order to species levels of taxonomic resolution (527 species; 77.6% of the taxa affiliated at the species level). Diet included insects of varying size (from 1mm to > 70mm), insects that emerge from water (e.g. Ephemeroptera, Odanata or Trichoptera), hard and soft-bodied insect species as well as spiders.

On one hand, we could not exclude that among the 527 prey species detected here, some may come from (i) environmental contaminants (Galan et al. 2018) or (ii) indirect predation (e.g. spider prey; Sheppard et al. 2005). The first possibility was likely to have only a minimal impact on our results due to the stringent filtering steps carried out before statistical analyses. However, it could not be completely avoided because for example, we are not able to filter the presence of coprophagous insect DNA contaminating guano before sample collection: in this case it comes out as prey despite the filters. The second possibility could not be distinguished from direct predation but should not be ignored in conservation perspectives. Prey indirectly consumed (*i.e*. prey of *R. ferrumequinum* prey) may exert a bottom-up control (Frederiksen et al. 2006), that is a direct impact on their predator populations (*R. ferrumequinum* prey), and therefore indirectly *R. ferrumequinum* itself. On the other hand, despite a high number of samples, our sampling effort was not high enough to describe the whole prey diversity of *R. ferrumequinum* for each sampling date and colony (except above the order taxonomic level of resolution). This was likely to be explained by the presence of highly numerous rare prey composing the secondary diet (300 taxa with only one occurrence in all the dataset) (Razgour et al. 2011; Vesterinen et al. 2013; Clare et al. 2009). This result suggested that the diversity of *R. ferrumequinum* diet should be even higher than described therein, and would require more samples to be fully estimated (Mata et al. 2018). Most bat diet studies are based on metabarcoding analyses of feces collected directly from captured bats (usually placed individually in cloth bags until they defecated), what limits the sample size per sampling session (usually < 30 samples; Vallejo et al. 2019; Aldasoro et al. 2019; Alberdi et al. 2020; Galan et al. 2018; Bohmann et al. 2018). Thus, using eDNA extracted from guano collected beneath colonies should be favored to describe bat diet as it enables to process a large number of samples while minimizing disturbance and stress for bats.

Beyond the description of a huge diversity of prey in *R. ferrumequinum* diet, our results emphasized the need to distinguish a core diet potentially constituting the foundation of *R. ferrumequinum* diet, and a secondary diet composed of numerous rare prey mainly occurring once in the samples over all colonies and sampling dates. The core diet was made of 15 common prey species shared by all the colonies, from medium to large size (10 to 40mm long for Coleoptera and Diptera, and 15 to 100mm width for Lepidoptera). A previous study based on microscopic analyses of *R. ferrumequinum* feces from England suggested that prey could be considered as ‘key prey’ or ‘secondary prey’ (Ransome et Priddis 2005). However, there was no clear definition of ‘key prey’ and ‘secondary prey’, and this classification was mainly based on prey abundance estimated from the percentage of prey volume in the feces. Considering different months throughout the summer, various moths, beetles, tipulids and ichneumonids were either identified as key prey or important secondary prey which became key prey when the previous key prey became scarce (Ransome et Priddis 2005). Most of the key prey described by Ransome and Priddis (2005) were also found in the core diet described therein (e.g. moth, beetles, tipulids), but some differences were observed. For example, we did not identify Geotrupes sp. and ichneumonids in the core diet, despite *Geotrupes spiniger* (36 occurrences) and nineteen ichneumonids taxa (from one to 12 occurrences) were detected in the secondary diet. Conversely, in the core diet we identified dipteran families (Limoniidae, Muscidae) which were not identified by Ransome and Priddis (2005). These differences between the two studies probably arose from the inability of the microscopic approach to detect and identify some prey of the core diet with precise taxonomic resolution, biasing the relative importance of each prey in *R. ferrumequinum* diet (Nielsen et al. 2018; Andriollo et al. 2019). Thus, our eDNA metabarcoding study based on the feces enabled to conceptualized more precisely the diet partition in core and secondary prey.

The existence of a core and a secondary diet suggests that *R. ferrumequinum* may exploit preferred taxa while simultaneously consuming on an opportunistic strategy a wide range of available prey. *R. ferrumequinum* ability to select prey relies on its particular echolocation system. This later accurately discriminates insects, their speed and trajectory while compensating for the Doppler shifts induced by its own flight – a unique adaptation of rhinolophid bats (von der Emde et Menne 1989; Ortega, Moreno-Santillán, et Zamora-Gutierrez 2016). Based on previous encounter experiences with prey, *R. ferrumequinum* is able to link prey-specific echo information with prey profitability, then to use it to inform hunting economic decisions from its perch (Koselj, Schnitzler, et Siemers 2011). The existence of a core and a secondary diet might therefore reflect the two hunting tactics used by *R. ferrumequinum*, that is prey searching in flight and prey searching from a perch (Jones et Rayner 1989). Because the first strategy induces a greater metabolic cost than the second one, *R. ferrumequinum* is expected to be more selective while perch-hunting (low cost - high yield tactic; Nadjafzadeh, Hofer, et Krone 2016) than while actively searching in flight (Koselj, Schnitzler, et Siemers 2011; Voigt et al. 2010). Thus, the core diet may be linked to perch-hunting and the secondary diet to flight-hunting. These two complementary foraging behaviors (selectivity and opportunism), that seem to drive the diversity and composition of the whole and core diets, are thus important processes enabling *R. ferrumequinum* dietary plasticity. While biological and ecological characteristics of *R. ferrumequinum* would favor such partition of its diet, it might not be unique to this bat species. Prey selection and the presence of numerous prey occurring only once in the diet have also been reported in other bat species, including *Myotis daubentonii* a water surface forager taking advantage of the acoustic mirror provided by water to detect its prey (Vesterinen et al. 2013; Vesterinen et al. 2016; Ciechanowski et al. 2007). To what extent the core/secondary diet partition can be generalized to insectivorous bat species and whether the echolocation system and hunting tactics of bats are associated with such diet partition remain important issues that need to be deeply addressed in the future using metabarcoding approach.

We detected strong spatio-temporal differences of the dietary composition. They were driven by the numerous rare prey of the secondary diet which were found only once in the whole sampling. This could reflect the importance of prey from the secondary diet to complement the core diet (Ransome et Priddis 2005). Indeed, the core diet might meet the energetic needs for development and reproduction whereas the secondary diet might play a role of diet completion to enable survival when essential prey are scarce (e.g. Mirhosseini, Hosseini, et Jalali 2015). The role of the secondary diet in energy completion is well known in human nutritional ecology (Taylor, Keim, et Gilmore 2005; Fanelli et Stevenhagen 1985; McGowan et al. 2012; Koehler, Harris, et Davis 1989) but remains unexplored in other animal species. For example, Caster (1980) showed that the core diet of women in a poverty population provided almost 70% of dietary energy but that the secondary diet provided important contribution to the average daily intake of proteins, iron, vitamins. Thus, further studies combining ecological and nutritional analyses will be of great interest to better assess the relative importance of a core and a secondary diet in providing energy and mineral intakes to insectivorous bats.

Our results suggested that prey availability, constrained by prey phenology, is an important factor influencing the diversity, composition and spatio-temporal variations of *R. ferrumequinum* diet. Differences in the availability of Diptera and Lepidoptera might influence the prey selection by *R. ferrumequinum* (Jones 1990), potentially to satisfy nutritional requirements (Nutritional wisdom hypothesis; Tracy et al. 2006). Indeed, we observed a shift from consumption of Diptera in July to Lepidoptera in August and Studier and Sevick (1992) showed that Diptera were globally richer in calcium, that is particularly useful for lactation (Barclay 1994; Kwiecinski, Krook, et Wimsatt 1987). The same shift was found by Jones (1990) and has also been described in other bat species (Clare et al. 2014).

The prey diversity peak of the whole diet in July might be explained by the variations of insect abundance over the summer, as a peak of insect diversity is often observed at or near the middle of this season (Wolda 1988). As such, the consumption of some of the most occurrent prey species identified as part of the core diet appeared to be clearly linked to their seasonal availability (e.g. *Triodia sylvina, Agrotis bigramma* and *Zeiraphera isertana*). However, if insect phenology was the main factor shaping *R. ferrumequinum* diet composition and diversity, we should not have observed such decrease of the core diet diversity in August. Indeed, most of the core prey were expected to fly at this moment (Table S2). Such discongruence corroborated the hypothesis that *R. ferrumequinum* dietary plasticity could be strongly influenced by other constraints. As we initially assumed, this temporal survey revealed important changes in both the diversity and composition of *R. ferrumequinum* core diet, that seemed to correlate with variations in female energetic and physiologic requirements during the maternity period. Indeed, the higher prey diversity observed in June and July suggested an increase of the niche breadth in response to high metabolic requirements and constraints during gestation and lactation. On the contrary, the lower diversity of prey observed in August could reflect a more selective foraging behaviour resulting from the decrease of maternity constraints but the still high energetic needs associated with pre-hibernation fat storage, migration to hibernacula and mating (Ciechanowski et al. 2010; Kunz, Wrazen, et Burnett 1998; Speakman et Rowland 1999). The ability to adjust foraging behavior with reproductive conditions, and hence energy requirements, has already been shown in bat species including *R. ferrumequinum* (Dietz et Kalko 2007). The decrease in flight manoeuvrability of pregnant bat females, as well as the decrease in flight distance and the increase in returns to the roosts of bat females during lactation period (Henry et al. 2002; Dietz et Kalko 2007) could dampen bat selectivity for particular prey during gestation and lactation and favor the ingestion of a larger spectrum of prey. This ingestion of potentially less profitable prey can be compensated by increased daily food intakes and higher digestion efficiency (e.g. changes in the gastrointestinal tract of adult females) (Reynolds et Kunz 2000; Encarnação et Dietz 2006; Kunz 1974). The ensuing decrease of diet diversity in late summer has already been observed for several bat species, for example *Plecotus auritus* (brown long-eared bat; Andriollo et al. 2019) and *Myotis lucifugus* (the little brown bat; Clare et al. 2014). All these bat species seemed to consume a greater volume of a limited number of prey taxa, what can lead to a decrease in diet diversity in late summer. As accumulating body reserves must be performed rapidly before the onset of winter and hibernation, the selective feeding strategy in August should target profitable prey species (e.g. rich in fatty acids; Levin et al. 2013; Krüger et al. 2014) and be associated with hormonal, metabolic and gut microbial composition/diversity changes (Kronfeld-Schor et al. 2000; Srivastava et Krishna 2008; Levin et al. 2013; Xiao et al. 2019). In other cases, such as the big brown bat *Eptesicus fuscus*, bats may exploit a wider variety of habitats and prey, what in turn results in an increase of diet diversity in late summer (Clare, Symondson, et Fenton 2014).

Our results also suggested that landscape features are important drivers of prey availability, and as such, influence the diversity, composition and spatio-temporal variations of *R. ferrumequinum* diet. Insect species richness and abundance are strongly influenced by landscape features such as plant species richness or landscape heterogeneity (Schuldt et al. 2019; Rundlöf et Smith 2006). Consistently, we showed that the more the landscape is differentiated between the colonies, the more their diet is differentiated and that the composition and diversity of *R. ferrumequinum* diet varied with landscape characteristics. For example, the lepidopteran *Laothoe populi* is known to be mainly associated with forests. As such it was much more detected in the two forested colonies (Beaumont and Annepont) than in the other ones. Conversely the coleopteran *Copris lunaris*, which is associated with dung in meadows, was less abundant in the two forested colonies. We also found that *R. ferrumequinum* diet diversity was lower in colonies surrounded by forests than by meadows and hedgerows. Thus the contrasting patterns of prey composition and diversity observed between the colonies dominated by forests and those by meadows and hedgerows are likely to reflect the adaptation of *R. ferrumequinum* to semi-cluttered foraging habitats (e.g. echolocation, hunting strategies; Dietz, Pir, et Hillen 2013; Jones et Rayner 1989). The large concentration and diversity of insects provided by hedgerows in agricultural fields should also enables the consumption of a wider variety of local prey (Verboom et Spoelstra 1999; Lewis 1969; Forman et Baudry 1984; Holland et Fahrig 2000). Therefore it is likely that these differences in prey communities associated with landscape features may lead to the selection of different profitable prey (Kolkert et al. 2020; Danks 2007; Clare et al. 2011).

At this point, we lack crucial information on the prey (availability and nutritional composition) throughout summer to validate the potential processes described above explaining dietary plasticity, and to evaluate the importance of selectivity in *R. ferrumequinum* foraging strategy (Koselj, Schnitzler, et Siemers 2011; Emlen 1966; Vesterinen et al. 2016; Tracy et al. 2006). Analyzing more deeply dietary variations between individuals from the same colony would also be particularly interesting. Indeed, as described for other bat and mammal species, local patterns might be influenced by the specialization of different individuals within colonies (Johnston et Fenton 2001; Thiemann et al. 2011; Bolnick et al. 2003). Differences in echolocation characteristics, flight and hunting performances between juveniles and adult bats may for example contribute to a wider spectrum of prey when young start to feed themselves compared to adults (Salsamendi et al. 2008; Rolseth, Koehler, et Barclay 1994; Arrizabalaga-Escudero et al. 2019). Genetic studies could help discriminating individuals and reconstructing pedigrees (Puechmaille, Mathy, et Petit 2007; Vesterinen et al. 2016). However, they would not give information on the reproductive status of female (non-reproductive, pregnant, lactating) nor on the age of bats (but see Wright et al. 2018 on non-degraded DNA); both would require bat captures.

### Conservation perspectives

At first sight, considering dietary plasticity as well as the wide spectrum of arthropods consumed by *R. ferrumequinum*, it might seem reasonable to consider that this bat species should not be at high risk when facing environmental changes affecting its prey distribution and abundance (Boyles et Storm 2007; Pratchett, Wilson, et Baird 2006; Twining et al. 2019; Owens et Dittman 2003). However, our results have emphasized the existence of a core diet - potentially essential for optimizing *R. ferrumequinum* fitness - which could be threatened by the modification of the landscape, the indirect effect of cattle anti-parasite drugs on the beetles and more globally the use of pesticides (Gonzalez-Tokman et al. 2017; Geiger et al. 2010; Pocock et Jennings 2008; Dietz, Pir, et Hillen 2013; Froidevaux, Broyles, et Jones 2019; Finch et al. 2020). Further studies are therefore needed to evaluate the effects of the core and secondary prey variations on bat life history traits and fitness. In particular, it is critical to assess whether these variations might significantly impact the demography and viability of *R. ferrumequinum* populations. This is all the more important given that it has recently been shown that some populations at the edge of *R. ferrumequinum* distribution might be at higher risk of extinction in the near future (Tournayre et al. 2019).

Besides, our results support a growing literature illustrating the significant role of insectivorous bats in arthropod pest control (Kolkert et al. 2020; Cohen et al. 2020; Aizpurua et al. 2018). Some large and/or chitinized pests had already been identified in previous *R. ferrumequinum* diet studies, such as the Coleoptera *Serica brunnea* and *Melolontha melolontha* or the Diptera *Tipula maxima* (Galan et al. 2018; Aldasoro et al. 2019; Jones 1990). However, no study had described as many deleterious or potentially deleterious insect species (one third of all occurrences) in *R. ferrumequinum* diet. Thus our results confirmed that *R. ferrumequinum* may be not only important as a sentinel of agricultural insect pests (chirosurveillance) but also as an efficient agent of pest control (Weier et al. 2019; Cohen et al. 2020; Maslo et al. 2017). This role could become even more important in the future because climate change is expected to favor the establishment and proliferation of many deleterious insects (Trumble et Butler 2009). Our results also support the presence of arthropods beneficial to agriculture (*i.e*. natural enemy of pests) in insectivorous bat diet (Cohen et al. 2020; Kolkert et al. 2020). Beneficial arthropods are rarely addressed in bat diet literature, probably because of the high scarcity of databases, but it is likely that insectivorous bats eat both pests and beneficial arthropods. Kolkert et al. (2020) showed that the diet of several Australian insectivorous bat comprised of around 1% relative abundance and richness of beneficial insects (predators, parasitoids, pollinators), emphasizing the service rather than the disservice these bats provide to agriculture. Assessing the impact of *R. ferrumequinum* on pest populations and determining which category of prey insectivorous bats eat, *i.e*. the relative quantities of pest and beneficial arthropods consumed, will be needed to further evaluate the effectiveness of *R. ferrumequinum* as agent of pest control.

## ACKNOWLEDGMENTS

We are very grateful to A. Cheron, J. Dechartre, M. Dorfiac, Y. Prioul and all the people involved in the guano sampling. We also would like to warmly thank E. Pierre for his help with arthropod pest species. Financial support was received from the LABEX ECOFECT (ANR-11-LABX-0048) of Université de Lyon, within the program “Investissements d’Avenir” (ANR-11-IDEX-0007) operated by the French National Research Agency (ANR). It also received support from the French National Research Institute for Agriculture, Food and Environment INRAE (EcoFA department), the internal funding from the CBGP laboratory, Nouvelle-Aquitaine region, Nouvelle-Aquitaine DREAL, and European Regional Development Fund as a part of a program driven by the Poitou-Charentes NGO. O. Tournayre PhD is funded by the LabEx CeMEB, an ANR “Investissements d’Avenir” program (ANR-10-LABX-04-01). Data used in this work were partly produced at the GenSeq platform through the genotyping and sequencing facilities of ISEM (Institut des Sciences de l’Evolution-Montpellier) and LabEx CeMEB. We are grateful to the genotoul bioinformatics platform Toulouse Midi-Pyrénées, and Sigenae group for providing help and computing resources thanks to Galaxy instance http://sigenae-workbench.toulouse.inra.fr.

## AUTHORS CONTRIBUTION

The study was conceived and designed by O.T., M.L., O.F.C., M.G., D.P. and N.C. The sampling schemes were designed by M.L., M.G., O.T., O.F.C., D.P., N.C., and conducted by M.L. Landscape data were extracted by D. P. Laboratory protocols were designed by M.G. and performed by M.G., O.T. and M.T. O.T., S.P., M.G. and M.T. performed the bioinformatics analyses and O.T. carried out the statistical analyses. A first draft of the manuscript was written by O.T, M.G., D.P. and N.C. All authors contributed to the writing of the final version of this paper.

## DATA ARCHIVING STATEMENT

Supplementary data deposited in Zenodo (10.5281/zenodo.3819911) include: (i) raw sequence reads of the five runs (fastq format), (ii) information on the samples, positive and negative controls, (iii) raw abundance table before and after filtering, (iv) final abundance table from the *R. ferrumequinum* samples only.

